# Seeing social: A neural signature for conscious perception of social interactions

**DOI:** 10.1101/2022.05.26.493596

**Authors:** Rekha S. Varrier, Emily S. Finn

## Abstract

Percepts of ambiguous information are subjective and depend on observers’ traits and mental states. Social information is some of the most ambiguous content we encounter in our daily lives, yet in experimental contexts, percepts of social interactions—i.e., whether an interaction is present and if so, the nature of that interaction—are often dichotomized as correct or incorrect based on experimenter-assigned labels. Here, we investigated the behavioral and neural correlates of conscious social perception using a large dataset in which neurotypical individuals viewed animations of geometric shapes during fMRI and indicated whether they perceived a social interaction or random motion. Critically, rather than experimenter-assigned labels, we used observers’ own reports of “Social” or “Non-social” to classify percepts and characterize brain activity, including leveraging a particularly ambiguous animation perceived as “Social” by some observers but “Non-social” by others to control for visual input. Observers were biased toward perceiving information as social (versus non-social), and activity across much of the brain was higher during animations ultimately perceived as social. Using “Unsure” reports, we identified several regions that responded parametrically to perceived socialness. Neural responses to social versus nonsocial content diverged early both in time and in the cortical hierarchy. Lastly, individuals with higher internalizing trait scores showed both a higher response bias towards social and an inverse relationship with activity in default-mode and limbic regions while scanning for social information. Findings underscore the subjective nature of social perception and the importance of using observer reports to study percepts of social interactions.

**Significance Statement:** Simple animations involving two or more geometric shapes have been used as a gold standard to understand social cognition and impairments thereof. Yet experimenter-assigned labels of what is social versus non-social are frequently used as a ground truth, despite the fact that percepts of such ambiguous social stimuli are highly subjective. Here, we used behavioral and fMRI data from a large sample of neurotypical individuals to show that participants’ responses reveal subtle behavioral biases, help us study neural responses to social content more precisely, and covary with internalizing trait scores. Our findings underscore the subjective nature of social perception and the importance of considering observer reports in studying its behavioral and neural dynamics.

## Introduction

A remarkable feature of human perception is how quickly and automatically we identify social information in the environment. This is exemplified by pareidolia, the phenomenon of seeing illusory faces in everyday objects (Liu et al., 2014; Palmer & Clifford, 2020); our sensitivity to body language and gaze directed at us (e.g., Mona Lisa effect; Todorović, 2006) and our tendency to overhear salient social cues in otherwise unattended information streams (e.g., cocktail party effect; Wood & Cowan, 1995).

In the brain, regions along the superior temporal sulcus (STS) have been classically associated with social cognition: the more posterior regions (pSTS) are involved in animacy perception (Lee et al., 2014; Sugiura et al., 2014) while the more anterior regions are involved in higher-level processes like mentalizing, language and gaze detection (Carlin et al., 2011; Deen et al., 2015). The recently proposed third visual stream (Pitcher & Ungerleider, 2021) posits a specialized pathway for processing social information that emphasizes the role of biological motion. This pathway proceeds from the primary visual cortex (V1) directly to the motion-processing region (V5/MT) followed by the pSTS and other STS subregions.

The association between motion and social perception is best exemplified by our tendency to spontaneously attribute social intentions to moving stimuli even when they consist of only simple geometric shapes (Bassili, 1976; Heider & Simmel, 1944; Scholl & Tremoulet, 2000). Whether such animations are seen as social depends largely on the movement patterns of the agents (Castelli et al., 2000; Gao et al., 2009). This phenomenon appears to transcend age (Gordon & Roemmele, 2014; Rochat et al., 1997) and culture (Barrett et al., 2005), although interestingly, is not found in monkeys (Schafroth et al., 2021). Individuals with certain neurological or psychiatric conditions— most notably autism—are less likely to perceive social interactions in these animations (Abell et al., 2000; Fong et al., 2017; Klin, 2000; Langdon et al., 2020) and show commensurately lower activity in typical social processing regions of the brain (Castelli, 2002; Herrington et al., 2007; Kana et al., 2009, 2015).

However, socio-perceptual variability is not limited to clinical populations. Neurotypical individuals also vary in if and how they perceive social interactions – even when animations are handcrafted by experimenters to be clearly social or non-social (Li et al., 2020; Nguyen et al., 2019; Rasmussen & Jiang, 2019). In neurotypicals, social perception covaries with traits like loneliness, anxiety, psychopathy, and autism-like phenotypes (Desai et al., 2019; Epley et al., 2008; Gardner et al., 2005; Kanai et al., 2012; Lessard & Juvonen, 2018; Powers et al., 2014; Sacco et al., 2016). Thus, using participants’ own percepts and individual trait scores will likely help us understand social perception better than experimenter-assigned labels. Here, we relied on participants’ responses rather than a “ground truth”. Further, because visual features are often not well controlled between handcrafted stimuli intended to be seen as social non-social, when possible, we also leveraged stimuli with similar or identical visual properties that nevertheless give rise to variable percepts across individuals.

In this study, we used a large dataset (n = 1049 healthy young adults) from the Human Connectome Project (Barch et al., 2013; Van Essen et al., 2013) to investigate the behavioral and neural correlates of conscious social perception. We found that compared to negative reports (“Non-social”), positive identifications (“Social”) were more frequent, faster and associated with less uncertainty, indicating a bias toward perceiving information as social. Occipital, temporal and prefrontal brain regions showed higher activity to “Social” information even when controlling for visual properties of animations. Some regions showed intermediate activity levels to “Unsure” reports, suggesting a parametric response to perceived socialness. Differences in activity between “Social” and “Non-social” percepts emerged early in time and in the cortical hierarchy. Both percepts and brain activity while viewing animations also correlated with internalizing traits. Overall, results paint a nuanced and individualized picture of social perception, suggesting that socialness is “in the eye of the beholder”.

## Materials and Methods

We primarily used data from the Social Cognition task of the Human Connectome Project (henceforth referred to as the “HCP study” or “HCP dataset”). The dataset is openly accessible, and consists of a large sample of neurotypical individuals, enabling us to study both the dominant and non-dominant percepts for specific animations. The social task was one of seven cognitive tasks that were run as part of the HCP task battery (Barch et al., 2013). In this task, participants watched ten 20s animations, of which five each were considered generally social and generally non-social (experimenter-assigned labels of Mental and Random, respectively). At the end of each animation, participants indicated whether they perceived a social interaction by pressing buttons (“Social”, “Non-social”, “Unsure”). To distinguish experimenter-assigned labels from observer responses, in this paper we use the terms Mental and Random for the former, and “Social”, “Unsure” and “Non-social” for the latter. In the HCP dataset, participants also completed trait-level questionnaires, which enable the study of inter-individual differences. Here, we focused on internalizing symptoms, which include anxiety, loneliness, and social withdrawal (details below in section *Correlation between traits, behavior, and neural activity*).

As participants had to wait until the end of each 20s-long animation to make a response, the behavioral data in the HCP does not reveal *when* the perceptual decisions were made, and any differences in decision time are likely to influence the trajectory of brain activity. Hence, we additionally performed an online study on 100 neurotypical individuals (henceforth referred to as the “online RT experiment”) to gain insight into when in the course of the animation-watching decisions might have been made, and how this varied across particular animations and individuals.

### Participants

This study used the Social Cognition Task dataset publicly available in the online HCP repository (https://db.humanconnectome.org/; for each participant, fMRI data sub-folders: *tfMRI_SOCIAL_RL* and *tfMRI_SOCIAL_LR*; behavioral: *\*TAB.txt*). Demographic information and trait scores used to study inter-individual differences were from the restricted category. We obtained complete fMRI data from 1049 individuals for the HCP Social Cognition task (ages 22-37; 562 female and 486 male).

For the online RT experiment that we conducted in July 2021, we recruited 100 neurotypical individuals (ages 18-48, mean = 23.2, SE = 0.64). from the United States and United Kingdom via the online platform Prolific (www.prolific.co, Palan & Schitter, 2018). Prior to the experiment, all participants read and acknowledged the virtual consent forms in accordance with the Institutional Review Board of Dartmouth College, Hanover, New Hampshire, USA.

### Stimuli

Stimuli in the HCP study were ten 20-second-long animations chosen from previous studies (Castelli et al., 2000; Wheatley et al., 2007). Longer animations had been snipped to 20s by the HCP researchers (Barch et al., 2013). The animations were presented in two runs with five animations each (run duration 3min 27s) interleaved with fixation blocks of 15s without jitter. The order of presentation was maintained across all participants (see Table 1). The number of Mental (M) and Random (R) animations were balanced within and between runs (run 1: 2M,3R; sequence M-R-R-M-R; run 2: 3M, 2R; sequence M-M-R-M-R. For a list of the animations as provided by the HCP and their properties, see Table 1. Note that in this paper, we drop the suffixes in the filenames (“−A” and “−B”) for convenience.

**Table 1:**
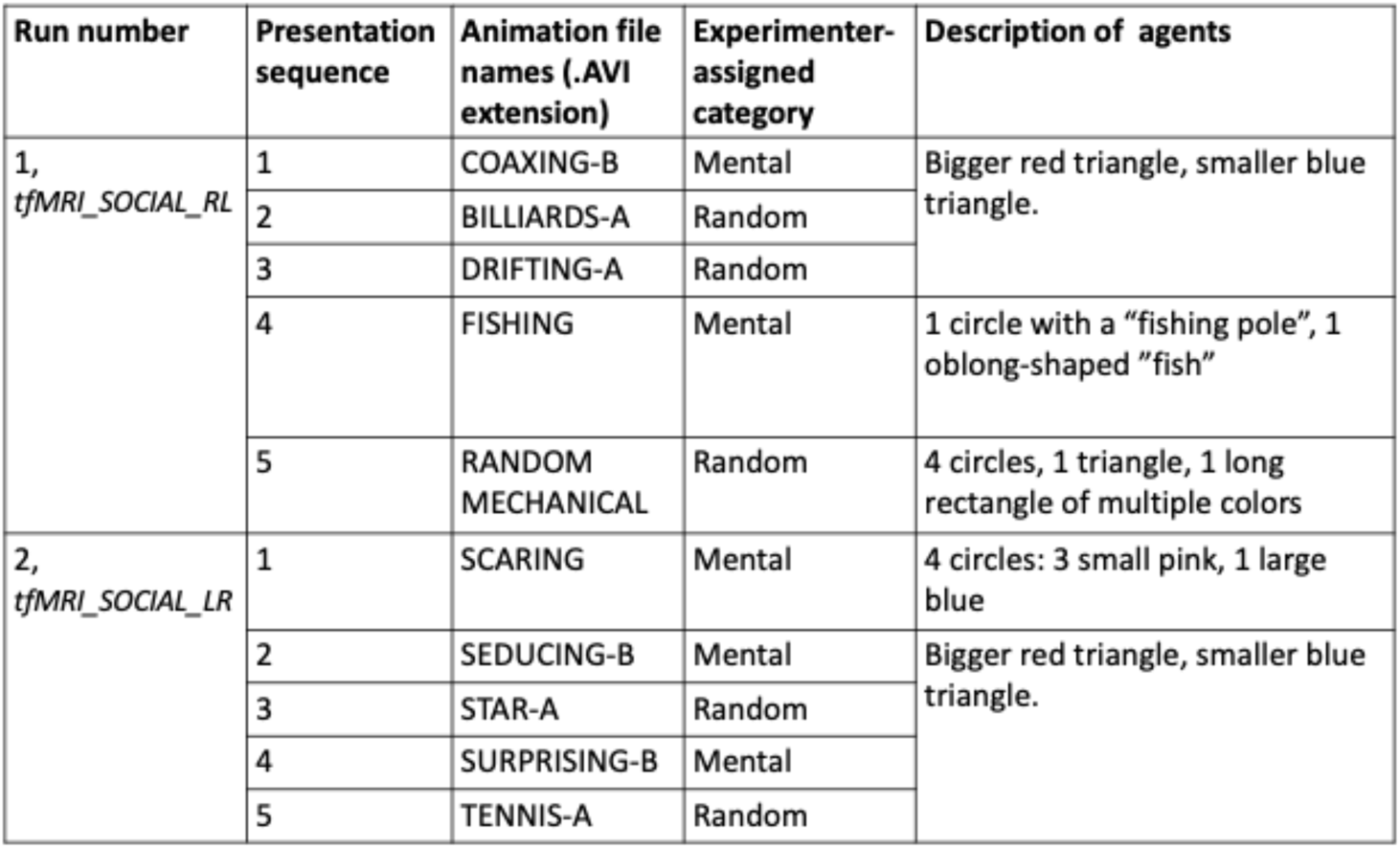
Animations used in the HCP Social Cognition Task.

Each animation consisted of two or more shapes in motion (“agents”) with or without stationary elements (“props”). Seven of them (3M, 4R) had a large red and a smaller blue triangle as agents, and the remaining three (FISHING, RANDOM MECH, and SCARING) were more diverse in the number, color, and form of agents and props.

For the online RT experiment, we presented the same animations used in the HCP study and in the same presentation sequence, with a self-timed break after the fifth stimulus in lieu of the break between the two runs in the HCP study. In the practice phase, we randomly showed either a generally social or non-social animation (that was not one of the 10 animations used in the main task) to each participant. For a social practice example, we used MOCKING-B from the HCP repository, and for a non-social practice example, we created a two-agent animation comparable in appearance to MOCKING-B using a custom app Psyanim (the latter available here: https://github.com/rvarrier/HCP_socialtask_analysis/tree/main/stimuli – link will be made public on publication. In the meantime, please get in touch with us for the file).

The differences in physical properties that we noted above amongst the HCP animations could have influenced both behavior and brain activity. Hence, we factored these into our data analyses steps either by comparing the brain activity for “Social” and “Non-social” responses *within* the same animation (i.e., same visual input) or by regressing out physical properties like the optic flow and mean brightness before comparing individual pairs of animations in the analysis comparing timecourses (explained in the sub-section *fMRI Timecourse Analysis* under *fMRI data analysis*).

The presence of these visual differences also motivated our decision to perform the online RT experiment to estimate decision times and consequently the selection of a pair of animations with similar decision times (details in the *fMRI data analysis* section). Lastly, to address the physical differences between specific animations, we also included animation as a grouping variable (“random effect”) in certain behavioral and fMRI data analyses when pooling data from multiple animations.

### Experimental design

In the HCP study, participants were given the following instructions about the task: “You will now watch short clips and decide if the shapes are having a mental interaction or not. For a mental interaction, press the button under your index finger. If you are not sure, press the button under your middle finger. For a random interaction, press the button under your ring finger. After each clip, there will be a response slide. Please respond while that slide is on the screen.” They had three seconds to respond. In our online RT experiment, participants were given similar instructions, but were asked to respond *twice* to each animation: once *during* the animation as soon as they made a decision (left/right arrows for “Social”/ “Non-social”) and a second time *at the end* of each animation within 3 seconds (left/right/down arrows for “Social”/”Non-social”/”Unsure” similar to the HCP study).

### Data acquisition and pre-processing

The fMRI data was acquired using a 3T Skyra scanner with 2mm isotropic voxels and a TR of 0.72s (see Barch et al, 2013 for more acquisition details). Each run comprised 274 scan volumes, and there were two runs per participant. We used minimally preprocessed voxel-wise fMRI data (Glasser et al., 2013), parcellated this into 268 parcels spanning the whole brain as per Shen atlas (Shen et al., 2013) and discarded the first five scan volumes (TRs) within each run to reduce initial artifacts. Next, to make BOLD response magnitudes comparable across participants, we z-scored parcel-wise timecourses in each run. Further, since our analyses were to be performed at the trial-level, we split the run time series into trial-wise timecourses of 40s each – i.e., 20s animations (28 TRs) flanked by 10s fixation periods (14 TRs) on either side (except for the first animation within each run which included only 6 pre-stimulus TRs). Data preprocessed in this manner was used for all fMRI analyses except one (the timecourse analysis, explained later) which required comparing two *individual* animations: COAXING and BILLIARD. Here, the z-normalization was done at the individual trial-level, to remove differences in mean activity that were due to the order of presentation (since order was not randomized between participants). In both cases, we lastly baseline-corrected each trial timecourse by subtracting the signal magnitude at the trial onset (i.e., from the TR immediately before stimulus onset).

In the online RT experiment, we excluded trials in which either of the two responses (“during” phase and “after” phase) were missing or where the two responses differed; the latter was done to ensure that the response time we see in the “during” phase correspond to the percepts reported in the end (to match the HCP task). Lastly, as a quality check, participants with fewer than 8 out of 10 good-quality (i.e., congruent) responses were also excluded, giving us 90 participants.

### Behavioral data analysis

We performed four analyses to measure whether there is a general bias toward social percepts, or in other words, a shift towards “Social” responses. For these analyses, we included only participants who responded to all 10 animations and in whom the response times (RT) were not unrealistically small (i.e., RTs < 100ms were excluded), giving an n=823 for these analyses. Our dependent variables were:

1. Percentages of “Social” and “Non-social” responses within participants; compared using a paired t-test
2. Decision criterion, the signal detection theory metric quantified as 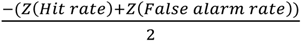 (Stanislaw & Todorov, 1999), where Hit rate and False alarm rate were computed for each participant as fractions of “Social” responses for animations labelled by the experimenters as Mental and Random, respectively.
3. Response time (RT) differences between “Social” vs. “Non-social” trials. We compared the RTs both using a non-parametric paired (Wilcoxon signed-rank) test and a more controlled linear mixed effects (LME) analyses to further account for the differences between individual animations. The LME model (LMEM) was of the form: *log*(*RT*) = *f*(*response*; *random intercepts: participant*, *animation*). The factor *response* was categorical with two levels: “Non-social” (coded as the base level) and “Social”, and analysis was performed using the Python package pymer4 (Jolly, 2018). We used the logarithm of the RT in seconds to bring the residuals of the LMEM closer to a normal distribution (which is an assumption for LMEMs).
4. Percentage of “Unsure” responses for the two animation labels (Mental, Random). These were compared using a logistic regression model: *uncertainty* = *f*(*stimLabel*; *random intercepts*: *participant*, *animation*) where the factor *stimLabel* was categorical [Mental, Random], and the dependent variable *uncertainty* had a value of “1” for “Unsure” response trials and “0” otherwise. Keeping Random (0) as the baseline in the analysis, positive/ negative regression coefficients for *stimLabel* would indicate a lower/ higher uncertainty in categorizing Random trials.

### fMRI data analysis

#### GLM-based regression

Our primary approach to fMRI data analysis was a general linear model (GLM) based on animation onset and offset. We computed the regression coefficients for each animation separately for the majority of analyses. For each animation, we fitted each parcel’s activity timecourse to a “slope” regressor (line steadily increasing from 0 to 1 from baseline to the duration of an animation, i.e., 20s, and padded by zeros before and after) that was convolved by the Glover HRF (Glover, 1999). (Preliminary analyses had indicated that a steadily increasing slope regressor captured more variance in the BOLD data than a traditional boxcar regressor.) This renders one slope regression coefficient (β) per parcel, participant, and trial (animation). We also performed a separate GLM analysis across all animations (details in the section below). For this analysis, we used a *run*-level regressor and estimated coefficients for each parcel, participant, and *run*. Similar to the slope regressors used at the trial level, regressor values increased (decreased) steadily during an animation labelled “Social” (“Non-social”) and were 0 at all other timepoints (including “Unsure” responses) – thus, the run-level regression coefficient here summarizes a *contrast* between “Social” and “Non-social”. For each participant, we then averaged these coefficients across the two runs.

#### “Social” vs. “Non-social”

To identify brain regions showing a consistent and generalizable difference between “Social” and “Non-social” responses, we compared the regression coefficients between “Social” and “Non-social” percepts in three analyses: (1) controlled for visual input, (2) controlled for decision times and (3) across all animations (Table 2). For analyses with individual animations, we included all participants who gave a valid response to the animation(s) in that analysis, resulting in slightly different numbers of participants in each analysis. Each analysis is described in detail below:

1. **Controlled for visual input**: We selected the most ambiguous animation, namely RANDOM MECH, since it has the relatively most balanced “Social” and “Non-social” response groups. We excluded participants who gave an “Unsure” response to this stimulus (leaving n=777) and then split regression coefficients based on observer responses (“Social”: n=107, “Non-social”: n=670, see Figure 1a), and compared them with two-sample t-tests assuming unequal variances.
2. **Control for decision time (COAXING vs. BILLIARD)**: We chose two animations which were most comparable in terms of the time taken to arrive at a decision about whether an animation was “Social” or “Non-social”. This was based on the data we obtained from the online RT experiment, where the decision time to report “Social” to COAXING (*median* = 3.45s, *SEM* = 0.27s) and “Non-social” to BILLIARD (*median* = 3.7s, *SEM* = 0.25s) were the closest and did not significantly differ (see Figure 2c and the results sub-section on decision time for more). Hence, we compared regression coefficients for each of these two animations within participants using a paired t-test. Note that we excluded participants who gave an uncertain or non-dominant response for one or both animations (i.e., who responded to COAXING as “Non-social” or “Unsure” or BILLIARDS as “Social” or “Unsure”), giving us n = 870 for this analysis.
3. **Across all animations (ALL):** We also performed a more general comparison between individuals’ responsiveness to “Social” vs. “Non-social” by identifying brain regions that show a mean run-level regression coefficient that is different from 0 (for details on how the run-wise regressor was estimated, see sub-section *GLM-based regression* above). To minimize biases due to missed responses, we only selected participants who had given all 10 responses and had complete fMRI data from both runs (n=814) using a one-sample t-test compared to 0.

**Figure 1.**
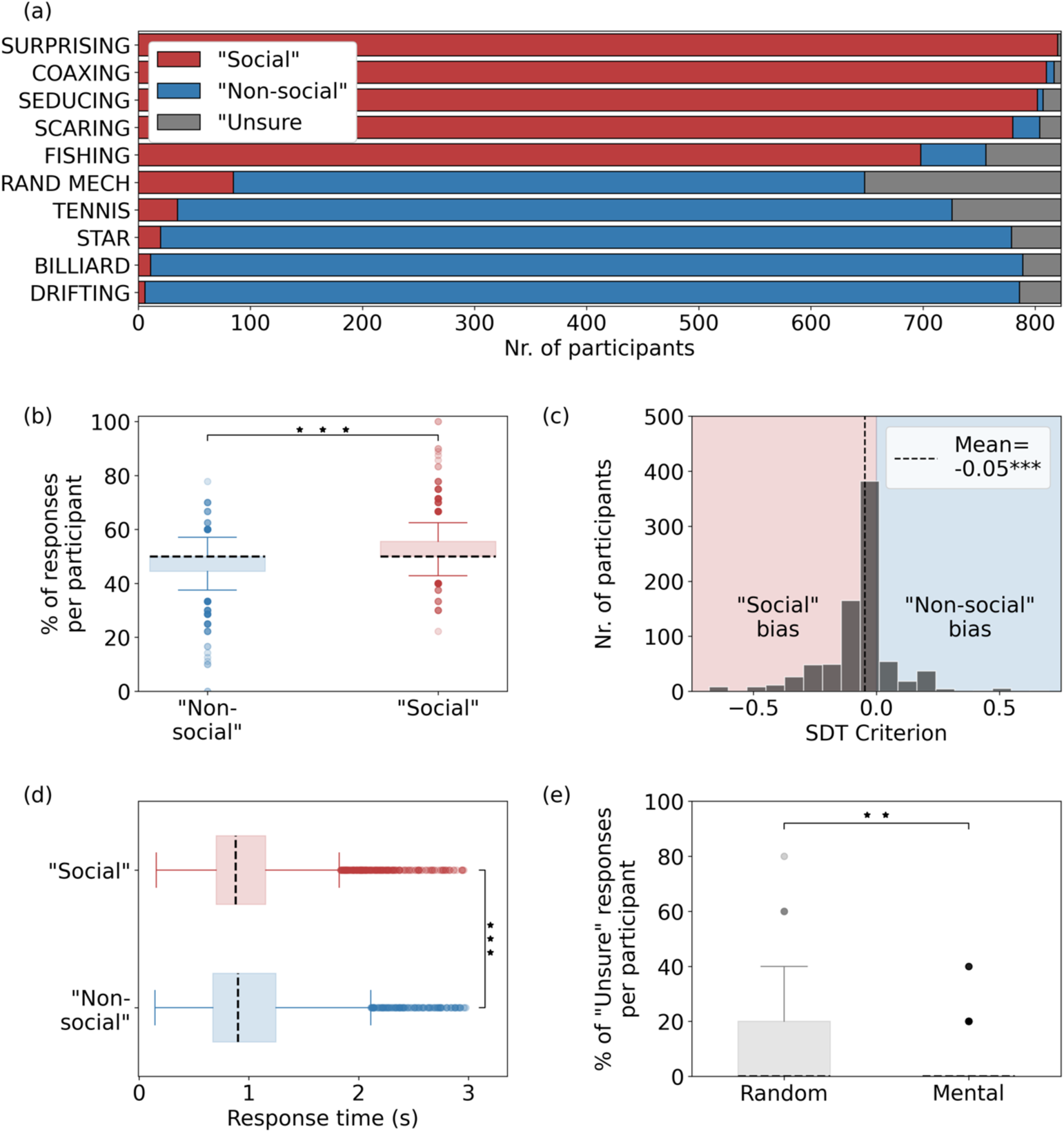
Behavioral data from the HCP participants (n=823) show a bias toward “Social” responses. (a) Number of responses per type (“Social”, “Non-social”, “Unsure”) and animation sorted from most to least “Social”. (b) Percentages of “Social” and “Non-social” responses. There was a higher number of “Social” responses (p < 10^−21^, paired t-test). (c) Signal detection theory metric “criterion” across participants based on experimenter-assigned labels. Mean criterion was negative (−0.05, p < 10^−17^, Wilcoxon signed rank test), indicating a bias toward false alarms (i.e., declaring an animation labeled Random by experimenters as “Social”). (d) Response time for “Social” and “Non-social” responses. “Social” responses tended to be quicker (Wilcoxon signed-rank, p < 10^−3^). (e) “Unsure” responses for animations labelled Mental and Random by experimenters. There was a higher percent of “Unsure” responses for Random responses (LMEM: Est. = −2.15, p < .005). **: p<.001, ***: p < .0001

**Figure 2:**
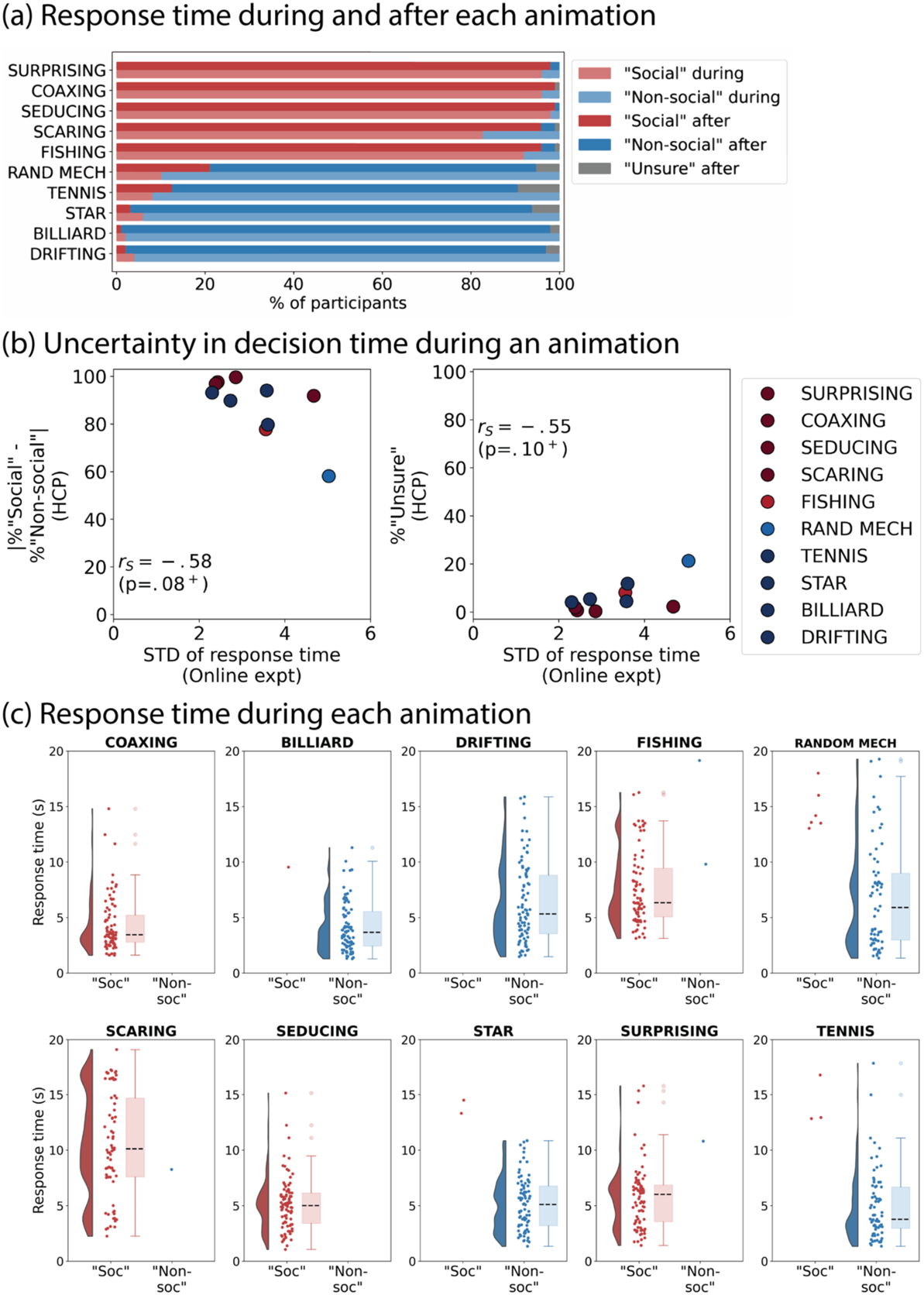
Results of the online RT experiment to characterize decision time for each animation. (a) Number of “Social”, “Non-social” and “Unsure” responses per animation made during (lighter shades) and after (darker shades) each animation. Order of animations on the Y-axis is the same as for the HCP data in Figure 1a. The degree to which animations were reported “Social” is comparable to the HCP behavioral data in Figure 1a. (b) Standard deviation of response time while watching each animation (in seconds; X-axis) vs. two indicators of uncertainty from the HCP behavioral data on the Y-axes (left: absolute difference between number of “Social” and “Non-social” responses, an indicator of how definitive responses for this animation were across participants; right: % of “Unsure” responses). Spearman (rank) correlation shows a trend (p ≤ .1, marked with ‘+’) for the animations with higher variation in response times in the online RT experiment (X-axes) to also a less definitive response (left) and a higher % of “Unsure” responses (right) in the HCP behavioral data. (c) Distribution of response times for “Social” and “Non-social” responses while watching each animation (in seconds). As seen in (b), decision times varied more for some animations than others. Note the similarity in the decision times between COAXING “Social” and BILLIARD “Non-social”.

**Table 2:** GLM analyses based on observer responses.

| Analysis Nr. | “Social” responder group | “Non-social” responder group | Rationale for the analysis | Statistical comparisons for fMRI | ID for the analysis used in figures and text |
| --- | --- | --- | --- | --- | --- |
| 1 | RANDOM MECHANICAL (within-animation, between-participant) |  | Controls for low-level input<br>Most ambiguous animation | Two-sample t-test | RANDOM MECH |
| 2 | COAXING | BILLIARD | Likely similar decision times *during* animation (as estimated from online RT experiment) | Paired t-test | COAXING-BILLIARD |
|  | (between-animation, within-participant) |  |  |  |  |
| 3 | All “Social” | All “Non-social” | Maximizes power by comparing all “Social” responses with all “Non-social” responses within participants | One-sample t-test after averaging run-wise estimates | ALL ANIMATIONS |
|  | Coded 1 (“Social”) and -1 (“Non-social”), in run-wise GLMs. |  |  |  |  |

Lastly, we identified brain regions that were significant in all three of the above comparisons and showed changes in the same direction (either “Social” > “Non-social” in all three comparisons or vice versa) at the FDR-corrected threshold (q < .05). We henceforth refer to this procedure as the “intersection analysis” and the resultant parcels as “social processing regions”.

#### “Social” vs. “Unsure” vs. “Non-social”

We also leveraged the “Unsure” responses to identify brain regions that responded parametrically to level of perceived socialness. We predicted that the neural response in such regions during animations ultimately marked “Unsure” would be intermediate to that of “Social” and “Non-social” responses. But note that intermediate does not necessarily mean halfway, and hence we performed conjunction analyses – i.e., we identified brain regions showing “Social” > “Unsure” and “Unsure” > “Non-social” (or vice-versa) and took the intersection of these. We performed this analysis across all the animations using an LMEM of the form: *beta = f(response*, *RI*: *participant)* which was performed separately for “Social” vs. “Unsure” (LMEM 1) and “Unsure” vs. “Non-social” (LMEM 2). In each LMEM, *response* was a categorical variable that has the values “Social” and “Unsure” in LMEM 1 (baseline “Unsure”), and “Unsure” and “Non-social” in LMEM 2 (baseline “Non-social”). Thus, a positive LMEM estimate for *response* would indicate a higher neural response corresponding to a higher perceived socialness. From this, we identified parcels which showed the same directionality for LMEM 1 and 2 at the multiple comparison-corrected threshold, and which were also in the set of social processing regions in the GLM analysis above.

#### fMRI Timecourse analysis

To identify the brain regions where the earliest differences in brain activity between “Social” and “Non-social” percepts emerged, we performed paired t-tests (within participant) for each timepoint (TR) between BOLD responses corresponding to a pair of “Social” and “Non-social” animations (COAXING and BILLIARD, respectively) in which decisions of whether the animation was social or non-social were likely made at comparable times *while* watching them as explained previously in the analysis sub-section *“Social” vs. “Non-social”*. To ensure that the differences between BOLD activity between COAXING and BILLIARD are not due to differences in basic visual input between the two animations, we performed these comparisons on the residual timecourses obtained after regressing out two low-level visual features, total optic flow and mean brightness. We first estimated these two features for each animation frame using the “pliers” package (McNamara et al., 2017), then down-sampled the resulting timecourses to match the temporal resolution of the fMRI data (i.e., the TR), z-transformed them and convolved them with an HRF. We then performed a linear regression on each participant’s trial timecourse (including 14TRs flanking the stimulus duration on either end like with the slope regressors described earlier) to regress out the changes in BOLD activity related to these features. We then used the resultant *residual* timecourses for COAXING and BILLIARD for the timecourse analysis. We compared these at each timepoint (TR) and for each parcel using paired t-tests (within participant). For each parcel, we thus identified the earliest timepoint at which BOLD activity begins to diverge. As additional consistency checks, we (1) only performed this analysis in the social processing regions that consistently differentiated between “Social” and “Non-social” in the GLM analyses, and (2) selected a TR *t* as the divergence point only if the difference between “Social” vs. “Non-social” at *t+1* was also significantly different in the same direction.

Note that this analysis does not factor in the hemodynamic lag. This is because although the HRF *peaks* a few seconds after an event (in our case, the animation onset), the neural responses to stimulus presentation should have begun instantly (Friston et al., 1994), so here we investigated where these earliest changes could be observed. Further, in using the median decision times from the online RT experiment for COAXING and BILLIARD as the expected decision time for the HCP dataset, we did not factor in the motor response delay (i.e., time taken after a decision has been made to press a button) in the online RT experiment. Hence it is possible that some of the pre-decisional processes closer to the decision time may have in fact been post-decisional. While we cannot exclude this possibility, this was unlikely since motor responses on arriving at a decision are typically quicker than the TR used in the HCP task (0.72s).

We also did not multiple comparison-correct across timepoints in this analysis since the primary goal was to identify the *earliest* differences in activity, and to infer this correctly, false negatives are less preferred to false positives. Further, in identifying the earliest timepoints, we only selected a region if the subsequent timepoint was also significant (p< .05 uncorrected), limiting the odds of a false positive further by 95%.

We also did not perform this analysis within the same animation (RANDOM MECH) and across all animations like in the GLM analysis (sub-section *“Social” vs. “Non-social”*) because of the heterogeneity in decision times both between reported percepts for the same animation and across animations (see Figure 2c). This means that the neural processes at each time point could have also been vastly different between “Social” and “Non-social” animations, thus making the comparison of timecourses less precise both within the same animation (RANDOM MECH) and across all animations.

### Correlations between traits, behavior, and neural activity

Past work has shown that individuals high on internalizing traits such as loneliness and anxiety tend to form illusory social connections by anthropomorphizing inanimate objects (Epley et al., 2008; Powers et al., 2014) and show smaller grey matter volumes in a brain region typically associated with social processing, the pSTS (Kanai et al., 2012). Here, we probed whether internalizing traits affect behavior and/or brain activity associated with social perception using the internalizing T-score provided by the HCP (Barch et al., 2013). This score is based on participants’ responses to the internalizing dimension questions which is part of the Achenbach Adult Self-Report questionnaire (ASR; Achenbach et al., 2017). Internalizing symptoms refer to symptoms like anxiety, depression, and withdrawal, and are typically contrasted with externalizing behaviors such as rule-breaking and aggression. The ASR was designed to assess behavioral, emotional, and social functioning across a wide spectrum of the population, so it is sensitive to individual differences (i.e., produces a range of scores) even in healthy/subclinical populations. We used the averaged T-scored participant-level internalizing score (labelled “ASR_Intn_T” in the HCP dataset; *M* = 48.72, *STD* =10.75, *range* = 30-97) for this analysis; see Figure 6a-c for the full distribution).

To assess whether internalizing score relates to a behavioral bias toward “Social” percepts, we correlated participants’ internalizing scores with the following behavioral variables using Spearman (rank) correlation: (1) the difference between % of “Social” and % of “Non-social” responses (calculated as percentages to control for missing data) and (2) the number of “Unsure” responses for Mental and Random trials, respectively. We also compared the internalizing scores between “Non-social” and “Social” or “Unsure” responders to the most ambiguous animation, RANDOM MECH. We tested the specificity of these correlations by additionally performing correlations with externalizing scores and comparing the two using the CorrelationStats package (https://github.com/psinger/CorrelationStats). To quantify if and where internalizing traits relate to brain activity while scanning animations for social information, for each parcel, we performed an LME analysis where the dependent variable was the slope regression coefficient, the fixed factor was internalizing score and the random factor was animation. This yields brain regions that respond proportionately to internalizing score in that individual across animations and parcels.

### Code availability

All the code for analyzing data from both the HCP and online RT experiment, as well as the anonymized data from the online RT experiment, will be made available upon publication here: https://github.com/rvarrier/HCP_socialtask_analysis. In the meantime, please get in touch with the authors for these.

## Results

In this study, we used behavior, fMRI data, and individual trait scores from the Human Connectome Project (HCP) social cognition task to characterize the behavioral and neural processes underlying conscious perception of social interactions. We started by evaluating the behavioral data for any response bias: are people more inclined to declare something “Social” (as opposed to “Non-social”)? We next identified brain regions that robustly differentiated between “Social” and “Non-social” percepts even when controlled for decision times and sensory information, including a subset of regions that showed a parametric response pattern to degrees of perceived socialness. Next, we used a timepoint-by-timepoint analysis to identify where and when brain activity begins to diverge between “Social” and “Non-social” percepts. Lastly, we studied the relationship between internalizing behavior scores, tendency toward social percepts, and brain activity while scanning for social information.

### Some animations are more ambiguous than others

First, we examined the degree to which participants’ percepts of “Social” versus “Non-social” information agreed with one another as well as the intended stimulus category. In the HCP social cognition task, participants passively watched ten 20-s animations of geometric shapes (Heider-Simmel-like; Castelli et al., 2000), see *Materials and Methods* sub-section *Stimuli* for a detailed description of the animations) and then made a behavioral response — “Social”, “Non-social” or “Unsure”—to indicate whether they perceived a social interaction in the animation. Five animations were intended to evoke social interactions (experimenter-assigned Mental) and five were not (experimenter-assigned Random). Although on average participants’ percepts aligned with experimenter labels, the degree to which animations were perceived as “Social” and “Non-social” varied considerably. This was true in both the HCP behavioral data and the secondary online dataset (online RT experiment) we collected to study the time taken for individuals to arrive at decisions while watching each animation (Figure 1a and 2a). While animations like DRIFTING and BILLIARD were seen almost unanimously as “Non-social”, animations like RANDOM MECH and FISHING had a higher percentage of the non-dominant percept as well as “Unsure” responses. This underscores the need to use participants’ own percepts to categorize what is or is not “Social” rather than experimenter-assigned labels. Further, in our analyses, we leverage this ambiguity by comparing neural activity corresponding to “Social” and “Non-social” responses within the most variably perceived animation (RANDOM MECH), thereby isolating activity associated with a conscious social percept while controlling for visual input.

### Responses are biased toward “Social”

Next, we used behaviorally reported percepts to determine whether there was a response bias towards “Social”. We hypothesized that evolutionarily, there may be a bias towards perceiving information as social, since the cost of a false positive (e.g., mistakenly thinking someone is trying to engage you in a social interaction) is lesser than that of a false negative (e.g., missing out on social cues that are important for group dynamics, reproduction, and survival). We predicted that this bias would manifest as a higher “Social” response rate, shorter response times for “Social” percepts, and more “Unsure” responses to animations labeled Random by experimenters (because of a reluctance to declare something entirely non-social). Our findings are described below:

1. **’Social’ responses are more frequent:** On comparing the frequency of percepts for each participant (limited to trials where participants were sure of their response—i.e., excluding “Unsure” trials), we observed that the percentage of “Social” responses was subtly but significantly higher (*M* = 52.89%, *SE* = 0.29%) than “Non-social” responses (*M* = 47.11%, *SE* = 0.29%; paired t-test, p < 10^−21^; Figure 1b).
2. **The response criterion further shows a bias towards “Social”:** Next, we computed criterion (c), a metric from signal detection theory that quantifies response biases. If the mean criterion c̅ is significantly different from zero, this suggests a bias in responses towards “Social” (c̅ < 0) or “Non-social” (c̅ > 0). We found that criterion was significantly negative (*M* = –0.047, *SE* = 0.006; Wilcoxon test p < 10^−17^; Figure 1c), further confirming the response bias towards “Social”. In this computation, we used the experimenter-assigned labels to show that although the experimenters aimed to create a balanced set of five Mental and five Random animations, actual observer reports indicate that individuals ended up perceiving more animations as “Social”. Thus, experimenter labels appear to be insufficient to explain all individuals’ “ground truth” percepts.
3. **Responders may have been quicker to declare something as “Social” than “Non-social**”: Next, to get at a more subconscious measure of perceptual decision-making for social information, we compared response times between “Social” (*med* = 0.87s, *SE* = 0.009s) and “Non-social” (*med* = 0.9s, *SE* = 0.012s) responses (Figure. 1d) and found that “Social” responses were overall faster (p < 10^−3^, Wilcoxon signed rank). Since response times could differ by animation due to their heterogeneity, we additionally performed an LME analysis with response (“Social” or “Non-social” [baseline]) as the fixed effect, and both animation and participant as random effects. We observed a trend towards shorter RTs for “Social” responses, but this did not reach significance (Est. = −0.037, p = .1).
4. **“Unsure” responses were more common for animations intended as Random compared to those intended as Mental:** We studied the distribution of “Unsure” responses between animations that were intended to be “Social” (Mental) or “Non-social” (Random) and noted that there was a higher percentage of “Unsure” responses in the animations intended as Random (*M* = 9.41%, *SE* = 0.5%; Figure 1e) compared to those intended as Mental (*M* = 2.70%, *SE* = 0.26%). This indicated that people were more reluctant to label something “Non-social” (as opposed to “Social”) when their confidence is low. In other words, they err on the side of false alarms rather than misses; this fits with the idea that misses are likely costlier than false alarms. We formally compared the frequency of “Unsure” responses using logistic regression with Mental (coded 1) and Random (coded 0) label as the fixed effect and participant ID and animation as random intercepts. Results showed higher uncertainty on Random trials even after accounting for the differences in animations (*Est*. = −1.61, *p* =.005).

To summarize, the behavioral data overall showed a bias towards “Social” responses based on percentage of each response type, response times and degree of uncertainty.

### Decision time as to whether an animation is “Social” varies widely between animations

In the HCP study, participants had to wait till the end of each animation (lasting 20s) to make a behavioral response. However, the decision as to whether an animation was “Social” or “Non-social” was presumably made sometime during passive viewing, although the decision time could have varied widely across animations and participants. This variability, in turn, might influence the timecourse of brain activity (e.g., visual attention for the same animation may be different when a participant makes a decision 2 seconds after the animation begins vs. 15 seconds after). Hence, getting information as to when decisions could likely have been made during each animation was critical to modeling and interpreting neuroimaging data. To this end, we performed an independent online behavioral study using the same animations where participants (final n = 90) were instructed to indicate their percepts as soon as they had arrived at a decision (“during” phase). To compare the results with the HCP study, participants were also instructed to respond at the end of each trial (“after” phase).

The consensus across participants of which animations were generally “Social” versus “Non-social” in the online sample was comparable to that of the HCP sample (see Figure 2a). As a corollary to this, the animations with high variability in decision times in the online RT experiment also tended to have less consensus across participants in the HCP study – the latter operationalized as (1) the absolute value of the difference between % “Social” and % “Non-social” animations (Figure 2b, left) and (2) higher number of “Unsure” responses (Figure 2b, right). The reaction time data from the “during” phase (Figure 2c) showed that while most responses were made in the earlier half of the 20 second animations, there was a high variability in decision time both within and across animations. This means that the brain activity corresponding to an especially ambiguous animation (e.g., SCARING, RANDOM MECH) could have been vastly different even amongst participants who reported the same percept for these, depending on when each participant made their decision and how this affected their attention before and after the decision. Hence, we identified two animations with the most comparable decision times, namely, COAXING (*med* = 3.45s, *SEM* = 0.27s), a predominantly “Social” animation, and BILLIARD (*med* = 3.7s, *SE* = 0.25s), a predominantly “Non-social” animation, whose decision times were not significantly different (Wilcoxon signed-rank test [paired], *T* = 1619, *p* = .57). We used this pair of animations in later analyses that required a control for decision time while watching an animation.

### Much of the brain responds more strongly to what is perceived as social information

In the next set of analyses spanning this and the next two sections, we used the fMRI data to understand where and when the brain distinguishes social from non-social information. For all fMRI analyses, whole-brain data were parcellated into 268 regions covering the cortex, subcortex, and cerebellum using the Shen atlas (Shen et al., 2013) to ease the computational burden of voxel-wise analyses.

In the first fMRI analysis, we focused on the question of “where” by comparing overall neural responsiveness while viewing animations ultimately deemed “Social” versus “Non-social”. In addition to regions along the STS which are known to be involved in animacy and interaction perception, we hypothesized that differences might emerge as early as visual regions. We compared “Social” and “Non-social” responses using a general linear model (GLM) approach— again, using the participant’s reported percept rather than the experimenter-assigned label as input to the model—in three separate contrasts to ensure results were robust to different confounding factors: 1) within the single most ambiguous animation (RANDOM MECH), which controls for visual input (since all participants saw the same animation, but reported different percepts; across-participants); 2) between two animations with similar decision times (COAXING vs BILLIARD), to control for effect of when the decision was likely made on the timecourse of brain activity during passive viewing (within-participants); and 3) across all ten animations, to maximize power and ensure generalizability (within-participants). We then identified social processing regions by taking the intersection of the regions showing a significant difference in all three analyses.

In total, 70 parcels showed “Social” > “Non-social” activity (FDR *q* < .05, black contours in Figure 3) consistently across all three comparisons, and no parcel showed “Non-social” > “Social” across analyses. Of these, 66 parcels showed positive activations for both the “Social” (β_”Social”_ > 0) and “Non-social” (β_”Non-Social”_ > 0) responses for both RANDOM MECH and COAXING-BILLIARD, suggesting that on the whole, much of the brain showed higher *activation* and not lower *deactivation* to “Social” compared to “Non-social”. These parcels spanned the occipitotemporal and prefrontal cortex, the cerebellum, and some sub-cortical regions (details in Table 3).

**Figure 3:**
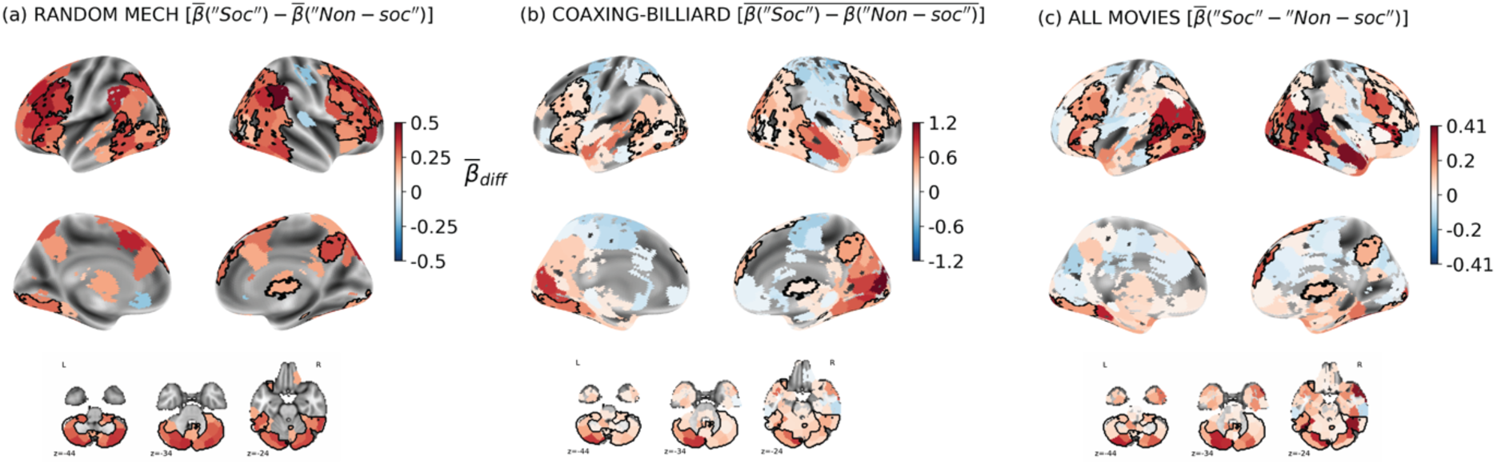
Identifying regions showing differential activity between “Social” and “Non-social” percepts. Mean differences between GLM regression coefficients (β) for (a) RANDOM MECH (mean (RANDOM MECH “Social”) – mean (RANDOM MECH “Non-social”), (b) COAXING-BILLIARD (mean(COAXING “Social”-BILLIARD “Non-social”)) and (c) ALL (estimated from run-level regressors, see Methods). Colored regions are significant at an uncorrected threshold (p < 0.05) in each of the three analyses, while black contours in a-c show the social processing regions significant after correction for multiple comparisons (FDR q < .05) in all three analyses. Note: Colorbar ranges are different between the three subplots, since each was estimated separately using different analyses, and hence the values shouldn’t be directly compared.

**Table 3:** Brain regions showing about “Social” v. “Non-social” differences.

| Lobe | Regions |
| --- | --- |
| Occipital | • Bilateral lateral occipital cortex<br>• Bilateral Occipital pole<br>• Bilateral occipital fusiform gyrus |
| Temporo-occipital | • Bilateral middle and inferior temporal gyrus (temporo-occipital part)<br>• Bilateral temporo-occipital fusiform cortex |
| Temporal | • Left posterior superior temporal gyrus<br>• Left posterior temporal fusiform cortex<br>• Left temporal pole |
| Parietal | • Right angular gyrus |
| Frontal | • Bilateral middle and inferior frontal gyrus<br>• Bilateral lateral precentral gyrus<br>• Bilateral frontal operculum<br>• Bilateral frontal pole<br>• Right superior frontal gyrus<br>• Parts of insula<br>• Left frontal orbital cortex |
| Sub-cortical | • Bilateral cerebellum (large parts of it)<br>• Bilateral brainstem<br>• Right thalamus |

### Some brain regions show parametric responses to degree of perceived socialness

The previous analysis identified social information-processing regions that robustly showed a higher response to information ultimately reported as “Social”. By leveraging “Unsure” responses as an intermediate level of perceived socialness between “Social” and “Non-social”, we further probed the neural correlates of conscious social perception—i.e., an “Unsure” response would indicate that some evidence for a social interaction was detected, but not enough to be fully confident in a “Social” response.

For this analysis, we probed which brain regions responded proportionately (quantified by the slope βs) to levels of perceived socialness — “Social”, “Unsure” and “Non-social”. Specifically, we sought to identify regions showing parametric responses, i.e., β_“Social”_ > β_“Unsure”_ > β_“Non-social”_ (condition S > U > NS) or β_“Social”_ < β_“Unsure”_ < β_“Non-social”_ (condition S < U < NS) using conjunction analyses across all animations (n=814) using separate LMEMs for “Social” vs. “Unsure” and “Unsure” vs. “Non-social” (see Methods for details). We further limited this analysis to the parcels that showed robust differences between “Social” and “Non-social” even when controlled for visual inputs and decision time (n = 70; cf. black contours in Figure 3).

38 parcels showed a consistent S > U > NS response pattern and of these, 35 survived multiple-comparison correction (q <.05) across all parcels (Figure 4a-b). This included posterior and inferior parts of the temporal cortex including parts of the motion-processing region V5/MT (with more parcels in the right hemisphere), middle and inferior frontal gyrus, precuneus, right thalamus and postero-lateral parts of the cerebellum.

**Figure 4.**
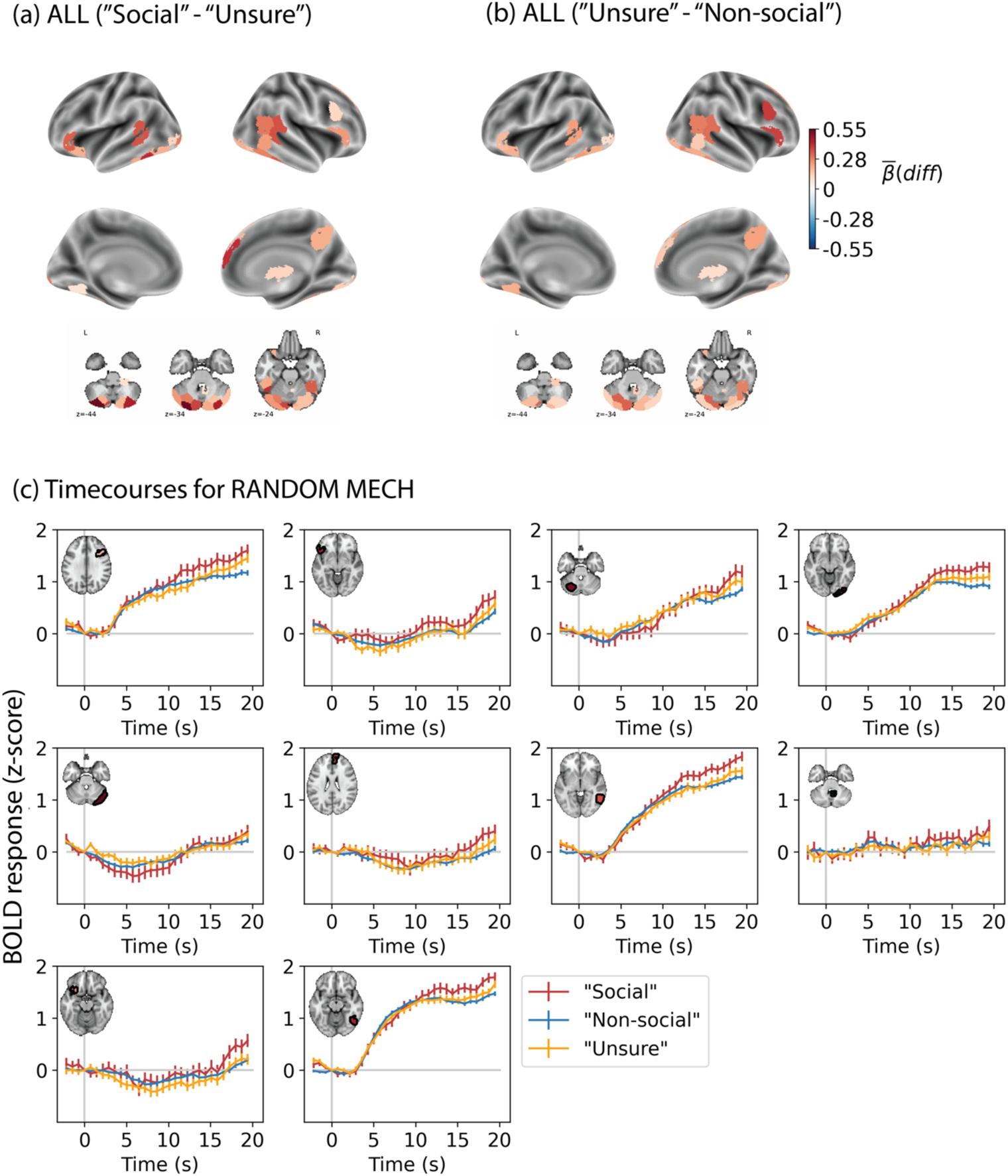
Brain regions showing parametric responses to social content. (a-b) Colored parcels show mean differences (FDR q < .05) in slope regression coefficients (“Social”-“Unsure” and “Unsure”-“Non-social”) for the 35 parcels which showed a graded response to perceived socialness (“Social” > “Unsure” > “Non-social” or vice-versa) across all animations and within the social processing regions obtained from the GLM analysis (cf. black contours in Figure 3). (c) Timecourses for the most ambiguous animation (RANDOM MECH) in 10 of the brain parcels sorted by how close “Unsure” is to the mid-point between “Social” and “Non-social” (details in text). These parcels included bilateral frontal regions, right middle and inferior occipito-temporal regions (including the anterior part of right V5/MT and parts of the cerebellum.

To see if similar trends emerge when controlling for visual input, we plotted the timecourses for each response type for a subset of the parcels showing parametric responses pattern to the most ambiguous animation (RANDOM MECH; Figure 4c). We plotted a sub-set of the parcels in which “Unsure” was the closest to the halfway point between “Social” and “Non-social” both in terms of the mean regression coefficient and the magnitude of activity at the end of the stimulus presentation period (20s) for each parcel and response (the rationale being that the signal during the final timepoints of the animation should be the best reflection of a participant’s ultimately reported percept). As expected, this results in brain regions which show parametric neural responses to degrees of reported socialness, albeit with large errorbars for the smaller groups (“Social” and “Unsure”).

Thus, it appears that there are at least some cortical and sub-cortical regions that show a graded response to degrees of social information. There was a higher number of such parcels in temporal, occipital and sub-cortical regions, although they were present across the cortex. This trend can be seen even on plotting the timecourses for the various responses to a single animation.

### Processing of social versus non-social information diverges early in time and across the cortex

The whole-trial-based analyses above showed that several regions spanning the whole brain are more responsive to information that is ultimately reported as social (versus non-social) even when controlling for decision time and visual input. However, this difference, especially in early visual regions, could reflect (1) the accumulation of evidence that *led* to the perception of an animation as “Social”, (2) the *consequence* of having perceived an animation as “Social” (i.e.., top-down attention effects on sensory regions), or (3) a combination of both. To gain a better understanding of the dynamics of evidence accumulation leading to a “Social” percept, we compared BOLD activity at each timepoint (TR) after stimulus onset to determine the timepoint of earliest divergence between “Social” and “Non-social” percepts.

To ensure that the differences observed at each timepoint are comparable in terms of the underlying cognitive processes (i.e., evidence accumulation versus decision-making versus post-decisional processes), we performed this analysis on the animation pair which likely had comparable decision times, namely COAXING-BILLIARD. Decision times for these animations were both early and close in time (as explained in *Materials and Methods* and the *Results* section for the online RT experiment, also see Figure 2c). These animations were similar visually with the same two triangular agents on the screen (see Table 1) – nevertheless, they did vary in the temporal dynamics and some low-level visual features. To minimize the effect on these on the BOLD activity, we regressed out the total optic flow and mean brightness from the BOLD responses of each animation and participant, and compared the residual COAXING and BILLIARD timecourses at each TR. To guard against spurious fluctuations early in the animations, we limited our analysis to the social processing regions (70 parcels) that showed a consistent difference in activity between “Social” and “Non-social” responses (Figure 3, black contours).

Differences in brain activity between “Social” and “Non-social” percepts emerged early, i.e., in TRs 1-3 after stimulus onset in many regions tested (Figure 5a). There were early differences between “Social” and “Non-social” in both hemispheres, both in posterior regions such as the fusiform gyrus, lateral occipital cortex, pSTS and posterior parts of the cerebellum as well as in frontal areas such as the lateral precentral gyrus, posterior parts of the middle and inferior frontal gyrus (MFG, IFG), the orbitofrontal cortex (OFC) in the left hemisphere, and the IFG and supplementary motor area (SMA) in the right hemisphere. Later TRs, which are more likely to reflect post-decisional activity, showed divergences in the bilateral inferior and superior frontal regions, the right precuneus, bilateral intraparietal sulcus (IPS) and bilateral posterior cerebellum. Within the regions that showed a significant overall differential activation for “Social”, the latest changes were seen in the left IPS and frontal pole and the right IFG and right anterior and medial cerebellum.

**Figure 5.**
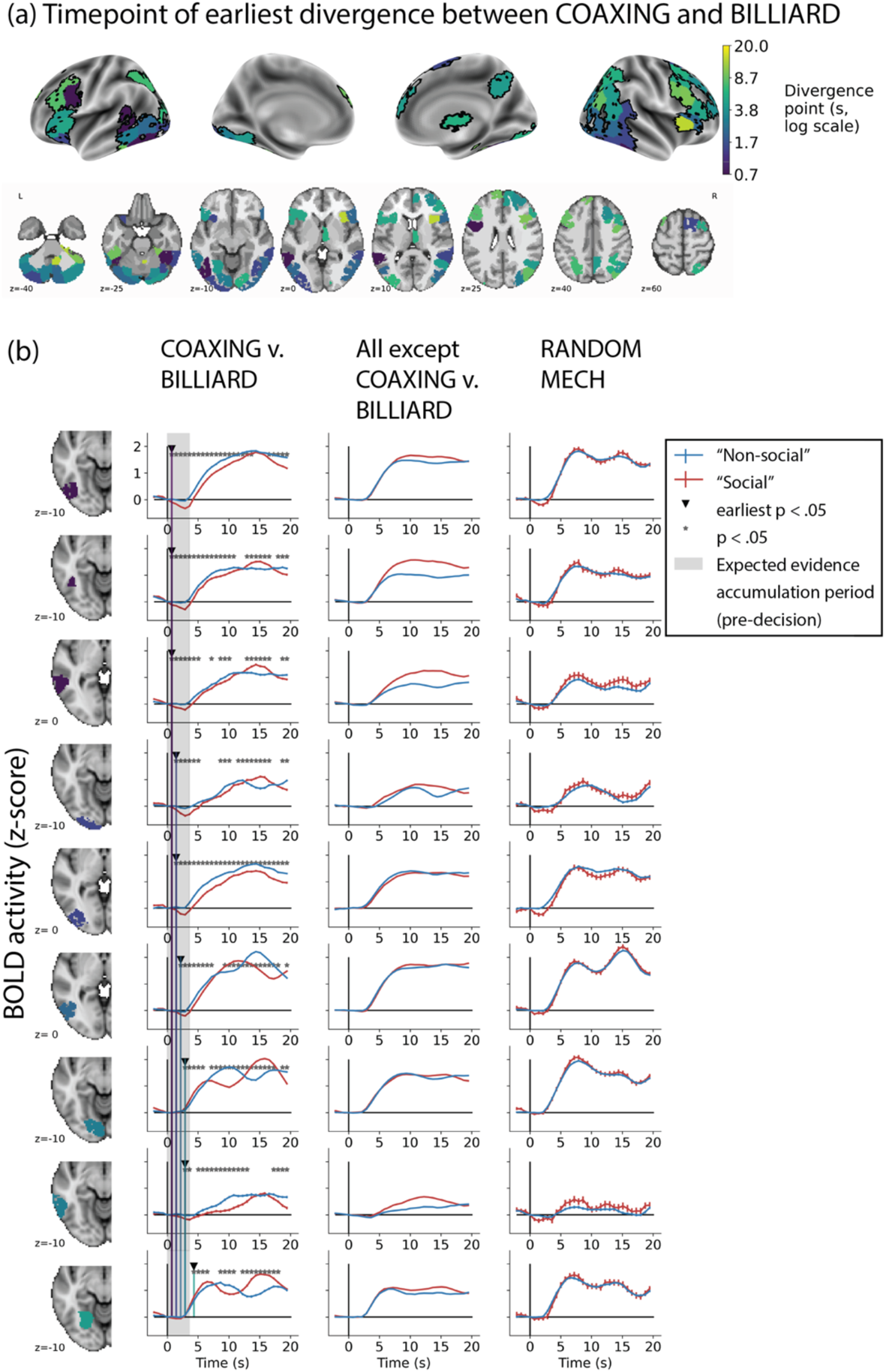
Timecourse analysis showing when and where differences between “Social” and “Non-social” percepts emerge. (a) Brain map of the earliest timepoint at which brain activity diverges between “Social” and “Non-social” responses for the COAXING and BILLIARD animations, respectively (within-participant analysis). Analysis was limited to the robust social processing brain regions (cf. Fig. 3, black contours), and BOLD signal timecourses were residualized with respect to the visual features of brightness and optic flow to minimize the effects of any differences in low-level sensory information between the two animations. Colors show how early (purple-blue) or late (yellow-green) activity diverged. (b) BOLD signal timecourses in the left posterior regions illustrating how “Social” and “Non-social” activity diverge in the pre-decisional period for COAXING and BILLIARD. Rows: regions are sorted by the earliest divergence TR and then from posterior to anterior. Columns: left, timecourses for the two animations matched for approximate decision time, COAXING (“Social”) and BILLIARD (“Non-social”), the main focus of this analysis. Others: timecourses from the same regions shown for two supporting analyses: across all animations except COAXING-BILLIARD (“Social” vs. “Non-social” response trials), middle; and for the most ambiguous animation, RANDOM MECH (“Social” vs. “Non-social” responders), right.

To visualize the earliest differences in the posterior regions and to understand how generalizable these dynamics are, we plotted (Figure 5b) the residual timecourses for COAXING-BILLIARD (left column, our main analysis) alongside the averaged “Social” and “Non-social” timecourses across all the other animations (All except COAXING-BILLIARD; middle column) and within the most ambiguous animation (RANDOM MECH; right column). The two latter analyses are not as well suited to pinpointing *when* differences emerged because decision times were likely more variable across individuals and responses for these animations (per our online RT experiment), thus making timecourses noisier and less comparable. Despite this, we see similar relative trends in these posterior regions (each row) as to when and how they distinguish between “Social” and “Non-social” reports. Responses emerged much later for the “All except COAXING-BILLIARD” condition in line with the later and more variable decision times for most animations; see Figure 2c). When comparing within the same animation (RANDOM MECH), we see trends emerging early on, although the magnitudes are smaller and the error for the “Social” responder group are large, possible because of the smaller group size (n = 107) compared to the majority percept of “Non-social” (n = 670). Note that the latter two timecourses are plotted only for visual examination and that we did not perform statistical analyses here.

To summarize, while watching an animation that was eventually reported as “Social”, differences in brain activity emerged early across much of the brain, involving both ventral visual processing regions and occipito-temporal regions involved in action and animacy detection as well as social cognition. The early jump in activity in these regions is in line with the recently suggested “third visual pathway” that projects directly from early visual cortex to the superior temporal sulcus and is specialized for social perception (Pitcher & Ungerleider, 2021).

### Individual differences in behavior and brain activity while viewing animations covary with internalizing symptoms

Lastly, we explored whether individual differences in behavioral and neural responses to social animations covaried with trait-level measures. Specifically, we focused on internalizing symptoms from the Achenbach Adult Self-Report Scale, because past work has shown that certain internalizing traits (e.g., loneliness, anxiety) are associated with a stronger tendency to perceive visual cues as socially salient. We hypothesized that individuals with higher internalizing scores would show stronger behavioral and neural reactivity to potentially social information.

Using the behavioral data, we tested whether the response bias towards “Social” (cf. Figure 1a) was even stronger for individuals higher on internalizing symptoms. Indeed, there was a positive relationship between the bias toward “Social” responses and internalizing score (Spearman rank correlation *r* =.104, *p* =.003, Figure 6a). We tested the specificity of this relationship by contrasting it to the correlation with externalizing trait scores, which index more “acting out” behaviors like rule-breaking and aggression and have not been linked to social perception tendencies. The correlation with externalizing symptoms was weaker and showed only trend-wise significance (*r*=.058, *p*=.096), and comparing the correlations showed a trend towards a significant difference between the two (*t* = 1.317, *p* = .094). Furthermore, individuals with higher internalizing scores were more likely to give a “Social” or “Unsure” (as opposed to “Non-social”) response to the most ambiguous animation, RANDOM MECH (“Social” or “Unsure”, *M* = 49.34, *SE* = 0.69; “Non-social”, *M* = 47.65, *SE* = 0.45; unpaired t-test, *t* = 2.054, *p* = .04, Figure 6b). Mean externalizing symptoms were also higher for the “Social” or “Unsure” group (*M* = 49.31, *SE* = 0.57) compared to the “Non-social” group (*M* = 47.98, *SE* = 0.38), although the difference was smaller (unpaired t-test, *t* =1.95, *p* =.05).

**Figure 6:**
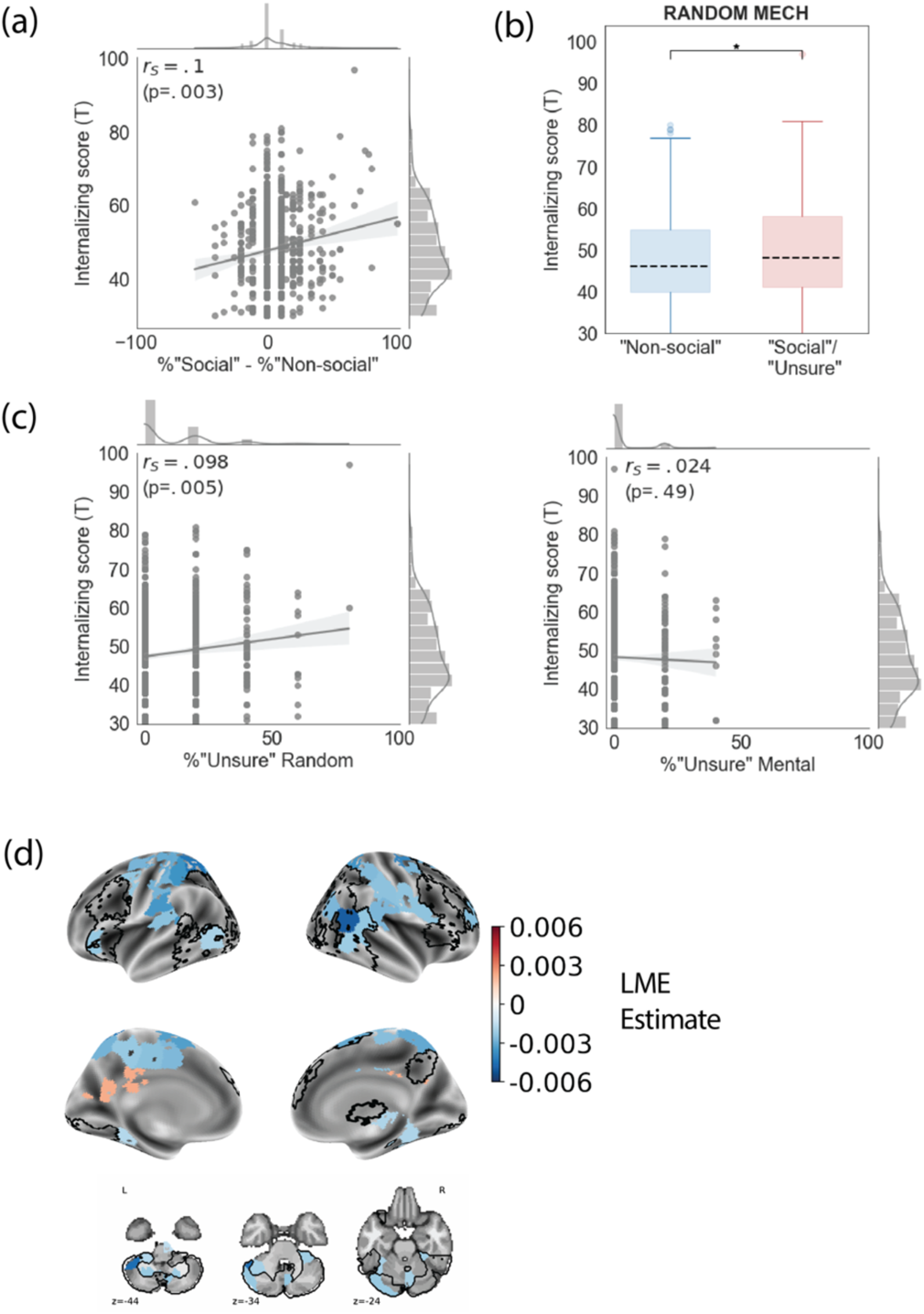
Relationship between internalizing trait scores, behavior and brain activity. (a) Response bias (% difference between “Social” and “Non-social” responses per participant) correlates positively with internalizing symptom score (Spearman correlation coefficient r_s_ = .1, p=.003). (b) Internalizing scores across individuals who perceived the most ambiguous animation, RANDOM MECH, as “Non-social” were lower than those for individuals who reported some degree of socialness to RANDOM MECH (“Social” or “Unsure” responses). * indicates p < .05. (c) Internalizing score correlates positively with the percent of “Unsure” responses per participant for the generally non-social animations (Random; left; Spearman r_s_ = .098, p =.005) but not for the generally social animations (Mental; right; Spearman r_s_ = .024, p = .49). These correlation magnitudes were significantly different (t= 2.47, p = .007). (d) LME estimates obtained by fitting the slope βs for each participant and animation to internalizing symptom scores per participant plotted over the brain. Colored parcels showed a significant relationship (FDR q < 0.05) and the social processing regions from the GLM analysis (cf. Figure 3) are shown in black. Most regions show a negative relationship with internalizing symptoms and there is only a partial overlap with the parcels that best differentiate “Social” and “Non-social” information.

Lastly, individuals with higher internalizing scores were also more likely to give an “Unsure” response to animations intended as Random (r=.098, p=.005), but not to animations intended as Mental (*r* = –.024, *p* = .49), indicating a preference for false alarms over misses when it comes to detecting social information (difference between correlations: *t* = 2.47, *p* = .007). Demonstrating specificity to internalizing symptoms, percent “Unsure” responses did not correlate with externalizing symptoms for either Random (Spearman *r* = .048, *p* = .17) or Mental (*r* = .01, *p* = .75) animations.

Together, these analyses support a link between internalizing symptoms and a greater tendency to perceive social information, perhaps driven by a homeostatic drive to seek social connections.

To understand whether neural activity while scanning for social information also covaried with internalizing symptoms, we related trial-wise regression coefficients to internalizing symptom scores in an LMEM (fixed effect: internalizing score; random effect: animation). In a whole-brain analysis, 50 parcels showed a significant relationship (*q* < .05, Figure 6d) between internalizing score and neural responsiveness. In 48 of these, the LME estimates were negative – i.e., as internalizing scores increased, the mean regression coefficients for an animation decreased – although all 48 parcels showed above-baseline activity as evidenced by the positive regression coefficients (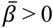 for all parcels). Thus, while individuals with higher internalizing scores showed positive activity in these regions when scanning animations for social information, the magnitude of this activity was lower than in individuals with lower internalizing scores. These relationships were seen in the right angular gyrus, the bilateral superior parietal lobule, supramarginal gyrus, regions along the dorsal midline, and left cerebellum (colored blue in Figure 6d). The two remaining parcels, left precuneus and posterior cingulate cortex (colored red in Figure 6d), showed a mean estimate that was positive but showed a net *de*activation in the group-level analysis (i.e., 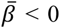), indicating that individuals with higher internalizing scores showed less deactivation in these regions. Thus, in most of the parcels that showed trait-dependent responses, the absolute magnitude of activity decreased with increasing internalizing symptom scores (which manifests as positive LME estimates when 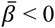 and negative LME estimates when 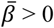).

Interestingly, the lateral occipital parcels from the social processing regions (shown as black contours in Figure 6d) were not as prominent here, showing only a partial overlap (13 parcels) with the parcels showing trait effects. In the overlapping parcels, which comprised bilateral occipito-temporal regions, the cerebellum and parts of the right superior frontal and left inferior frontal gyrus, individuals high on internalizing traits showed overall less reactivity in many brain regions while scanning the environment for social interactions. To reconcile this decrease in neural reactivity (Figure 6d) with the observed increase in behavioral sensitivity (Figure 6a-c), one interpretation is that these individuals have a lower threshold for the amount of neural activity required to declare something “Social”.

## Discussion

In this study, we investigated behavioral and neural signatures of social signal detection using a large dataset of neurotypical young adults. Behavioral responses showed a small but consistent bias toward perceiving information as social (as opposed to non-social), which manifested as a higher number of “Social” responses and a reluctance to report information as “Non-social”. We used the observers’ own labels of what was social and non-social to then identify brain regions that differentiate conscious social percepts, controlling for both visual input (RANDOM MECH) and decision time (COAXING-BILLIARD), and found that widespread patterns of brain activity differentiate conscious social percepts. A few brain regions also showed parametric responses to degrees of perceived socialness (“Social” > “Unsure” > “Non-social” responses). We further noted that brain activity for information ultimately deemed “Social” diverged from “Non-social” both early in time and in the cortical hierarchy. Lastly, we found that a trait-level measure of internalizing symptoms (e.g., loneliness, anxiety) could explain some of the variability in percepts and brain activity, such that individuals with higher internalizing traits had a higher tendency to perceive information as social yet lower reactivity in neural systems while scanning for this information.

Humans have been described as an “obligate social” species, evolutionarily tuned to social interactions (Rutherford & Kuhlmeier, 2013). In our results, both the bias towards “Social” responses in the behavioral data and the covariation between internalizing symptoms and sensitivity to social signals, which could reflect a homeostatic drive to seek social connection (Tomova et al., 2020), are in line with this. Moreover, previous studies have shown that people who report greater loneliness tend to form illusory social connections (Epley et al., 2008), overattribute animacy even in the absence of clear humanlike features (Powers et al., 2014) and have greater attention and memory for social cues (Gardner et al., 2005).

Past fMRI studies of animacy and social interaction perception using stripped-down geometric shape animations have primarily used two types of stimuli: short animations with simple, controlled motion profiles (e.g., Blakemore et al., 2003; Lee et al., 2014; Schultz et al., 2005; Tavares et al., 2008) or complex, scripted animations (Castelli et al., 2000; Nguyen et al., 2019; Osaka et al., 2012). Both sets of studies have primarily identified bilateral pSTS as relevant to intentional motion processing together with the lateral occipital cortex (LOC), angular gyrus, superior parietal lobule and medial prefrontal cortex. In this study, we observed differences between social and non-social percepts in these regions but additionally in the occipital pole, the left temporal pole and dorsolateral and ventrolateral prefrontal cortex. Several of these regions also showed parametric responses to degrees of perceived socialness. We did not observe a strong right-hemisphere dominance in this study as in some past work (Lee et al., 2014; Pitcher & Ungerleider, 2021). These disparities could be the result of the higher sensitivity we get as a result of the large sample size in the HCP compared to most other studies (n well below 100) and/or the use of observers’ own responses as stimulus labels instead of experimenter-assigned categories. The latter could be the bigger reason, since several of these additional regions were not seen even in past studies of the same HCP dataset (Barch et al., 2013; Li et al., 2020; Westfall et al., 2017). Future users of the HCP social task dataset are hence cautioned to not rely on experimenter-assigned labels alone.

In the timecourse analysis, we observed that the brain starts responding differently to social information early in time across postero-lateral visual processing regions, even before participants had likely arrived at a decision about whether an animation was social. Activity in these regions may therefore reflect pre-decision evidence accumulation processes, in which participants are using visual cues to determine whether agents are moving in a manner consistent with an intentional social interaction. Of note, these differences were unlikely to be due to differences in visual inputs since we regressed out the optic flow and brightness from each timecourse prior to the analysis. Early differences also emerged in the pSTS—an area critical to the third visual stream hypothesis (Pitcher & Ungerleider, 2021)—and lateral precentral gyrus, OFC and SMA. The presence of early differences in frontal regions involving the OFC is in line with previous accounts of a coarse-to-fine visual processing, where an early coarse information processing wave works in parallel with the slower more detailed processing via the ventral stream in spontaneous perception (Bar et al., 2006; Baror & He, 2021).

Brain activity while scanning animations for social information was lower for individuals with high internalizing scores in several regions including parts of the default mode network (angular gyrus, precuneus and posterior cingulate cortex) – some of which have been linked to depression, early-onset psychosis and anxiety (Collin et al., 2021; Nair et al., 2020; Sheline et al., 2009; Zhao et al., 2007) – and some of the social information processing regions (occipito-temporal, frontal, cerebellar) that robustly emerged from the group-level GLM analysis. Together with the positive correlation between the “Social” bias in behavior and the internalizing scores, this suggests that individuals with higher internalizing scores may have a lower threshold for declaring something “Social”.

One limitation of this dataset is the heterogeneity of the animations, which were taken from two studies (Castelli et al., 2000; Wheatley et al., 2007) with vastly different visual features. Paradoxically these animations are also not optimal to study ambiguous perception since they *did* have a dominant percept. Nevertheless, the large sample size enabled us to study percept-level as well as inter-individual differences. Further, the order of animations was not counterbalanced. Even though the timecourse analysis controlled for key visual differences, we acknowledge that we used two specific stimuli presented in a certain sequence in the HCP task, and hence cannot confidently extend these results to *all* social and non-social stimuli. We therefore propose this as an early step to future studies that investigate social processing pathways using more controlled stimuli.

Another limitation is that in this task, individuals had only three response options. However, participants’ confidence levels in these responses may have been vastly different, especially since the data showed that some animations were more ambiguous than others. Further, even given a response of “Social”, the nature of the particular interaction perceived could have varied across individuals, even for the same animation. This could have reduced the effect sizes and can be overcome in future experiments by using richer behavioral characterizations of percepts (e.g., continuous response scales) and/or indirect physiological measures like pupillary responses or electromyography.

In summary, we describe behavioral and neural processes that underlie how people arrive at conscious percepts of social information. We find evidence that neurotypical individuals are primed to detect social signals, and that this detection process is reflected in widespread brain activity that happens early in time and in the cortical hierarchy. We also find considerable heterogeneity among individuals’ percepts of particularly ambiguous information – i.e., information that may or may not be social in nature – and describe one trait-level factor that may influence behavioral and neural tendencies toward social versus non-social percepts. Together, results indicate the need for a more nuanced view of social perception in which socialness is in the “eye of the beholder”.

## Conflict of interest statement

The authors declare no competing financial interests.

## Acknowledgments

This project was supported by a NARSAD Young Investigator Award (grant number 28392) from the Brain & Behavior Research Foundation and by a Neukom CompX Faculty Grant from the Neukom Institute for Computational Science at Dartmouth College.

## Notes

### Competing Interest Statement

The authors have declared no competing interest.

